# PHD1-dependent hydroxylation of RepoMan (CDCA2) on P604 modulates the control of mitotic progression

**DOI:** 10.1101/2025.05.06.652400

**Authors:** Jimena Druker, Hao Jiang, Dilem Shakir, Fraser Child, Vanesa Alvarez, Melpomeni Platani, Andrea Corno, Constance Alabert, Adrian T. Saurin, Jason R. Swedlow, Sonia Rocha, Angus I. Lamond

## Abstract

Prolyl-hydroxylases (PHDs) are oxygen sensing enzymes that mediate the hydroxylation of proline residues. In mammals, three PHD isoforms (PHD1-3) are responsible for proline hydroxylation of Hypoxia Inducible Factor (HIF) alpha, a key regulator of the hypoxia response. In the accompanying paper (Jiang et. al., 2025) we report development of a mass spectrometry-based method to reliably identify proline hydroxylation (OH-Pro) sites on proteins and use this to identify a PHD-dependent OH-Pro modification at Pro604 on the protein RepoMan (CDCA2), a regulatory subunit for protein phosphatase PP1γ, with important roles in mitotic progression and cell viability.

Here, we investigate the functional significance of hydroxylation of RepoMan at P604. During M phase, the PP1-RepoMan complex dephosphorylates Thr3 of Histone H3 (H3T3) on chromosomes arms to ensure the correct localisation of the chromosomal passenger complex (CPC) at centromeres. We show that siRNA depletion of PHD1, but not PHD2, increases H3T3 phosphorylation in prometaphase-arrested cells. In cells depleted of endogenous RepoMan, exogenous expression of wild type RepoMan, but not a RepoMan P604A mutant, restored normal H3T3 phosphorylation localisation in prometaphase arrested cells. RepoMan P604 is located proximal to the Short Linear Motifs (SLiMs) that function as binding sites for the serine/threonine Protein Phosphatase 2A (PP2A). The interaction of RepoMan and PP2A-B56γ is reduced in cells expressing RepoMan P604A. Moreover, analyses in both fixed and live cells released from a prometaphase arrest, show that expression of the RepoMan P604A mutant delays completion of mitosis, results in defects in chromosome alignment and segregation and increases levels of cell death. These data support a role for PHD1-mediated prolyl hydroxylation in controlling progression through mitosis, acting, at least in part, via hydroxylation of RepoMan at P604 regulating the interaction of RepoMan with PP2A during chromosome alignment and thereby controlling the levels of Histone H3 phosphorylation at Thr3.

## Introduction

Proline hydroxylases (PHDs) are part of a large family of 2-oxoglutarate, iron and oxygen dependent enzymes (2-OGD) ^1^. PHDs are best known for their role in hydroxylating and controlling Hypoxia Inducible Factor (HIF) levels in normal oxygen, iron and 2-oxoglutarate conditions ^2^. They are also sensitive to metabolites analogous to 2-oxoglutarate. including succinate and fumarate ^3,4^, oncometabolites, such as L2-hydroxyglutarate ^4^ and certain amino acids ^5^. PHDs are thus intricately linked to oxygen and metabolic sensing and control ^1^.

There are three PHD enzymes in mammals, termed PHD1, PHD2 and PHD3, all of which have HIF hydroxylating functions ^6^. However, PHD2 is the dominant PHD for HIF hydroxylation ^7^, with PHD1 and PHD3 being required for negative feedback loops ^8,9^. PHD1 is not hypoxia inducible. Hydroxylation of HIFα results in the formation of a high affinity binding site for the von Hippel Lindau (VHL) protein, which is the recognition component of an E3-ubiquitin ligase, targeting HIFα for rapid degradation by the proteosome ^10–13^. Given the importance of PHDs in controlling HIF levels, several PHD inhibitors have been developed by the pharmaceutical industry ^14^, and these are now approved for use in human patients suffering from anaemia derived from chronic kidney disease ^14^.

Identification of additional PHD targets, outside the HIF family, has been surrounded by controversy over the last 15 years, confounded by the fact that only PHD2 has a significant phenotype in knockout mice ^15^. However, recent studies have provided evidence, based upon a combination of genetic, biochemical, molecular and cellular biological data, identifying additional targets for all PHD enzymes ^16^. When analysing the function of these novel targets, broadly, PHD2 targets seem to relate to signalling and metabolism, while PHD1 targets align to gene regulation and cell cycle ^16,17^.

We have developed a robust, mass spectrometry-based method for the enrichment and identification of proline hydroxylation sites on target proteins, allowing the detection of PHD targets in cell and tissue extracts. In the accompanying manuscript ^18^, we validate this MS approach and use it to identify a set of putative PHD target proteins. We have previously shown that PHD1 contributes to the control of cell cycle progression ^19,20^. Interestingly, one of the novel PHD1 target proteins identified by Jiang et al. ^18^, was CDCA2, also known as RepoMan, which is a cell cycle regulated protein phosphatase 1 (PP1) interacting protein^21^ RepoMan functions as a regulatory subunit for protein phosphatase PP1γ, with important roles in mitotic progression, chromosome architecture and cell viability^21,22^. RepoMan localisation to chromatin is regulated during the cell cycle by phosphorylation and dephosphorylation mechanisms and this is required for proper chromosomal passenger complex (CPC) localization and mitosis progression ^23,24^.

Here, we characterise the functional significance of RepoMan hydroxylation at P604, as identified by Jiang et al.^18^. We demonstrate that RepoMan hydroxylation at P604 is sensitive to PHD inhibitors, PHD1 depletion and increased levels of fumarate. RepoMan P604 hydroxylation is required for efficient mitotic progression from prometaphase into metaphase with loss of P604 hydroxylation causing delayed mitosis, defects in chromosomes alignment and segregation and cell death. These results support a critical role for PHD1 in controlling cell cycle progression and expand the repertoire of validated protein targets whose function depends upon site-specific proline hydroxylation.

## Results

### RepoMan interacts with PHD1 in asynchronous cells

Previous studies have shown that RepoMan localises to chromatin during interphase but rapidly dissociates from chromatin and becomes diffusely distributed when cells enter mitosis. Upon anaphase onset, RepoMan loads back on to chromatin and remains there until the next mitosis ^21,23,24^. First, therefore, we confirmed by immunofluorescence that we could reproduce the previously reported localization behaviour of endogenous RepoMan throughout the cell cycle, both in fixed HeLa cells (Figure 1A) and in live HeLa cells using YFP-RepoMan (Figure 1B). These results support the previous conclusions and show that RepoMan loads onto chromatin in HeLa cells at anaphase onset and remains associated with chromatin throughout the subsequent interphase, until is dissociates from chromatin when cells enter the next mitosis.

**Figure 1:**
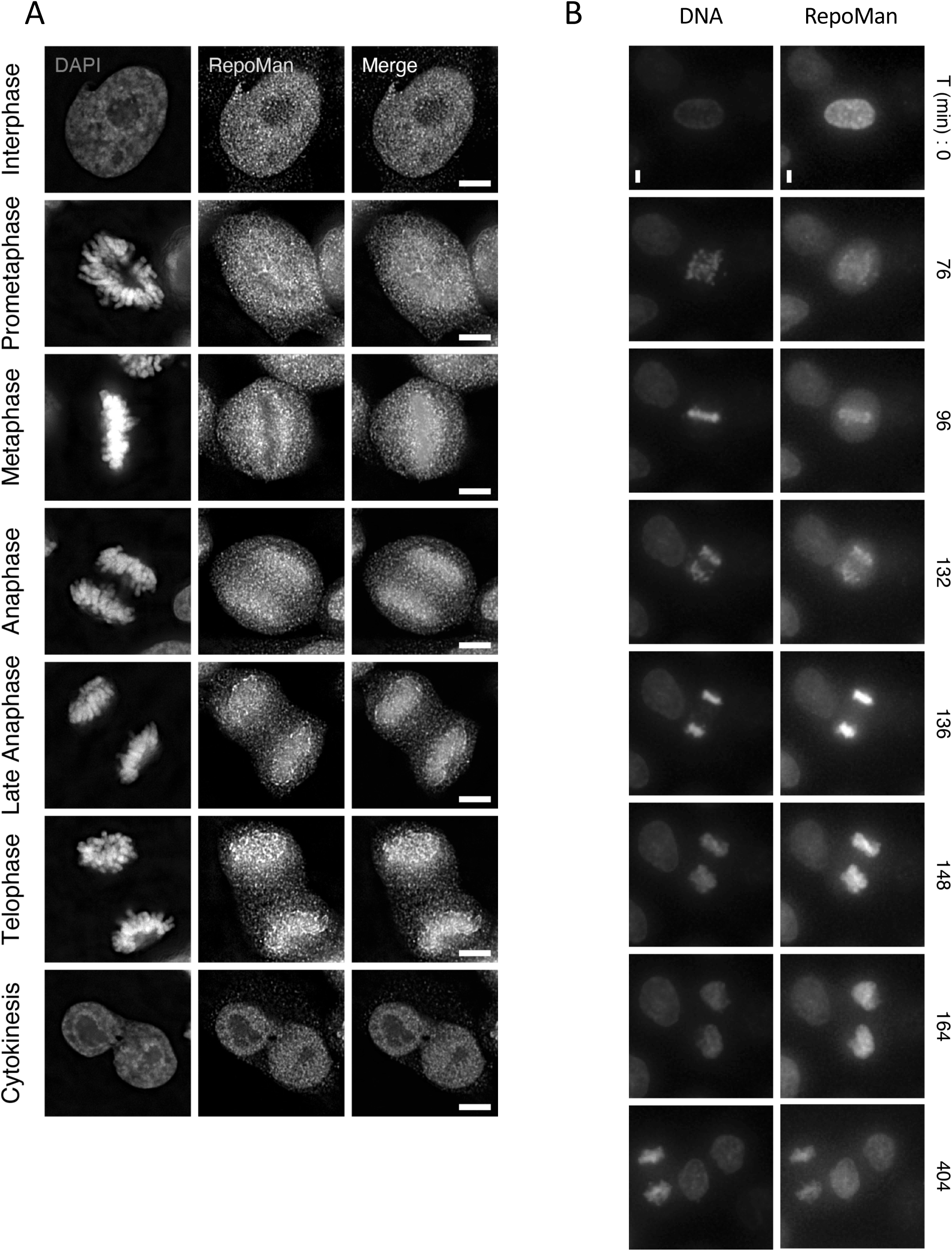
RepoMan localisation across the cell cycle. (A) Immunofluorescence analysis of endogenous RepoMan in HeLa cells along the cell cycle. CDCA2 antibody was used to detect endogenous RepoMan. DNA was stained with DAPI. Scale bar 5µm. (B) Time lapse of YPF-RepoMan in cells treated with thymidine for 24h and release in normal media. DNA was stained with sirDNA. YFP-RepoMan in Green and DNA in Red Scale bar 5µm.

Next, we compared the localisation of RepoMan and PHD1 across the cell cycle in asynchronous cells by analysing the localisation of endogenous RepoMan in HeLa cells transiently expressing EGFP-PHD1, both during interphase and mitosis (Figure 2A). EGFP-PHD1 localised in nuclei during interphase, consistent with previous results showing PHD1 to be bound to promoter regions and involved in hydroxylation of histone H3 ^25^. RepoMan also localised to nuclei during interphase but became enriched around the chromosome periphery during prometaphase (Figure 2A). However, at anaphase onset, RepoMan, as expected, loads on to chromatin, but PHD1 remains diffuse around the segregating anaphase chromosomes (Figure 2A). These results show that the localisation patterns of PHD1 and RepoMan are not identical, consistent with PHD1 acting on multiple different target proteins and not only RepoMan. The main overlap in localisation occurs during interphase and (pro)metaphase, assessed by Pearson correlation ^26^, potentially allowing enzyme and substrate to colocalize and interact (Figure 2B).

**Figure 2:**
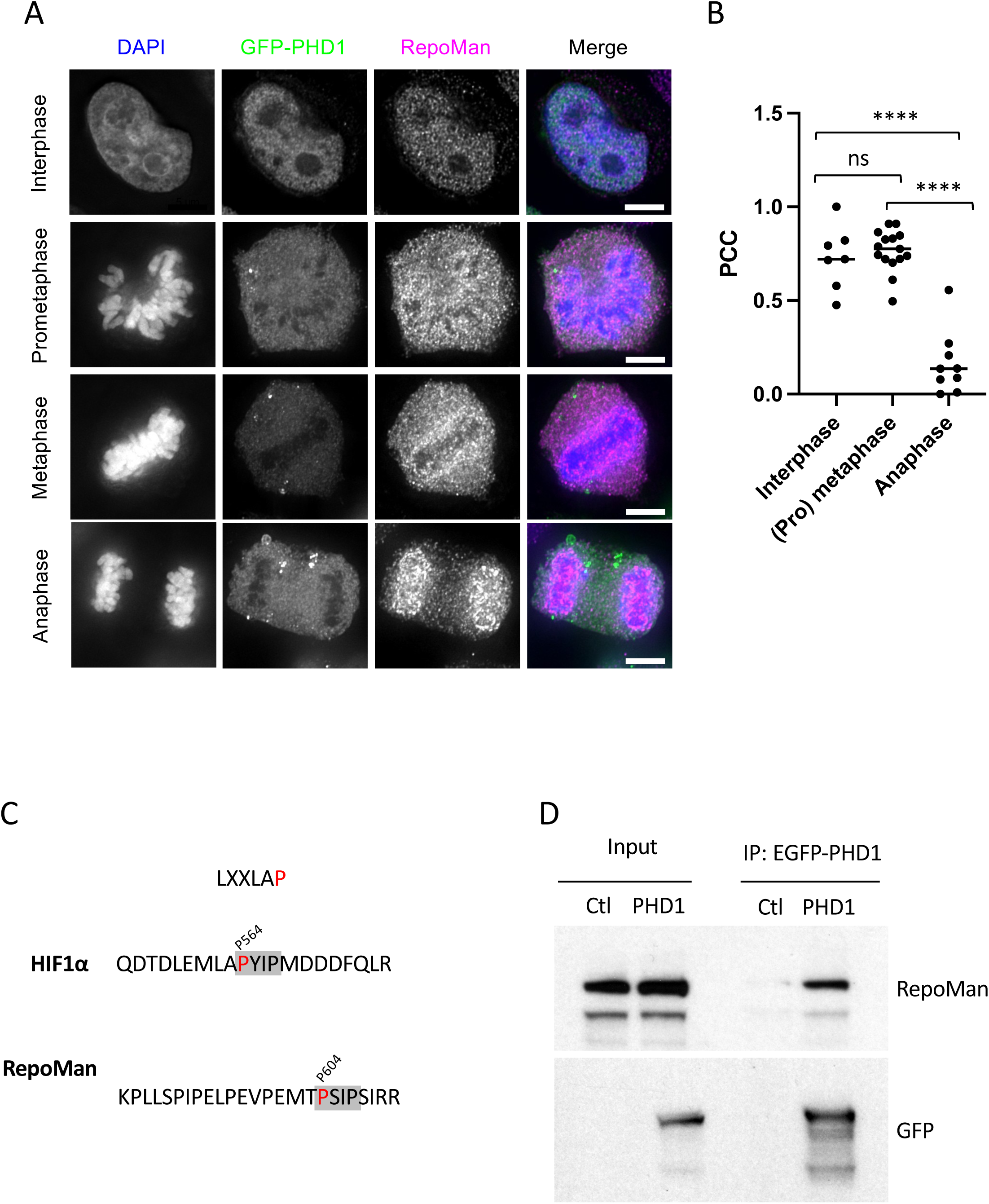
Endogenous RepoMan interacts with EGFP-PHD1. (A) Asynchronous HeLa cells were transiently transfected with 1µg of EGFP-PHD1 expression vector and after 48h were fixed with 4% PFA and subjected to immunofluorescence using CDCA2 and GFP antibodies (endogenous RepoMan and EGFP-PHD1 respectively). DNA was stained with DAPI. The Images show the localization of endogenous RepoMan and EGFP-PHD1 along the cell cycle: interphase, prometaphase, metaphase, and anaphase. Scale bars represent 5 µm. (B) Colocalization of GFP-PHD1 with endogenous RepoMan is shown. The graph displays the median of Pearson’s correlation coefficient measured (using Costes’ automatic threshold) in asynchronous cells captured in different stages of the cell cycle (Interphase, (pro)metaphase and anaphase). Unpair t test p= 0.0531 interphase vs (pro) metaphase, p < 0.0001 Anaphase vs (pro) metaphase, p < 0.0001 Anaphase vs Interphase. (C) Short sequence similarity between HIF1α and RepoMan around prolines 564 and 604 (PYIP, PSIP) respectively. (D) Co-immunoprecipitation between EGFP-PHD1 and endogenous RepoMan. Asynchronous HeLa cells were transiently transfected with EGFP-PHD1. After 48h cells were lysated and subject to immunoprecipitation using GFP trap magnetic beads and analysing by western blot.

RepoMan does not possess the consensus motif for proline hydroxylation (LXXLAP) present in HIFα. However, a short sequence next to Pro604 in RepoMan (PSIP), is similar to the PYIP sequence next to Pro564 in HIF1α (Figure 2C). These amino acids are involved in the PHD2-HIF1 oxygen-dependent degradation domain interaction through hydrogen bonds and hydrophobic interaction with residues in PHD2 that are conserved in PHD1 ^27^ and may thus be involved in the RepoMan-PHD interaction we detect in cells. To test this further, we analysed the interaction between RepoMan and ectopically expressed EGFP-PHD1 in asynchronous cultures of HeLa cells. Co-IP experiments, using GFP-Trap, show that endogenous RepoMan is in a complex with EGFP-PHD1 (Figure 2D). Together, these results are consistent with a model in which RepoMan and PHD1 can potentially interact throughout the cell cycle, except during anaphase, suggesting that the hydroxylation of RepoMan P604 by PHD1 might occur before chromosome segregation.

### Reduction in either PHD1 levels or activity increases H3T3ph on chromosome arms

RepoMan functions as a regulatory subunit for protein phosphatase PP1γ, with important roles in mitotic progression and cell viability (Trinkle-Mulcahy, 2006). During prometaphase and metaphase, PP1-associated RepoMan dephosphorylates Thr3 of Histone H3 (H3T3) on chromosomes arms ^28^. This prevents the recruitment of the chromosomal passenger complex (CPC) and promotes the correct localisation of the CPC complex to centromeres, where H3T3 serves as a docking site (Figure 3A). Proper localisation of the CPC is crucial to ensure correct chromosome alignment and segregation during mitosis ^23,24,28,29^. Reduction of RepoMan expression by siRNA has been reported to increase H3T3 phosphorylation on chromosome arms in prometaphase cells ^23,24,28,29^. To determine if this result could be replicated in our experiments, RepoMan was knocked down in HeLa cells using siRNA and H3T3 phosphorylation analysed in nocodazole-arrested cells, both by immunofluorescence (Figure 3B) and Western blot (Figure 3C). This confirmed the previous reports that RepoMan depletion coincides with increased H3T3 phosphorylation. Importantly, this result validates the use of H3T3 phosphorylation as a readout of RepoMan function.

**Figure 3:**
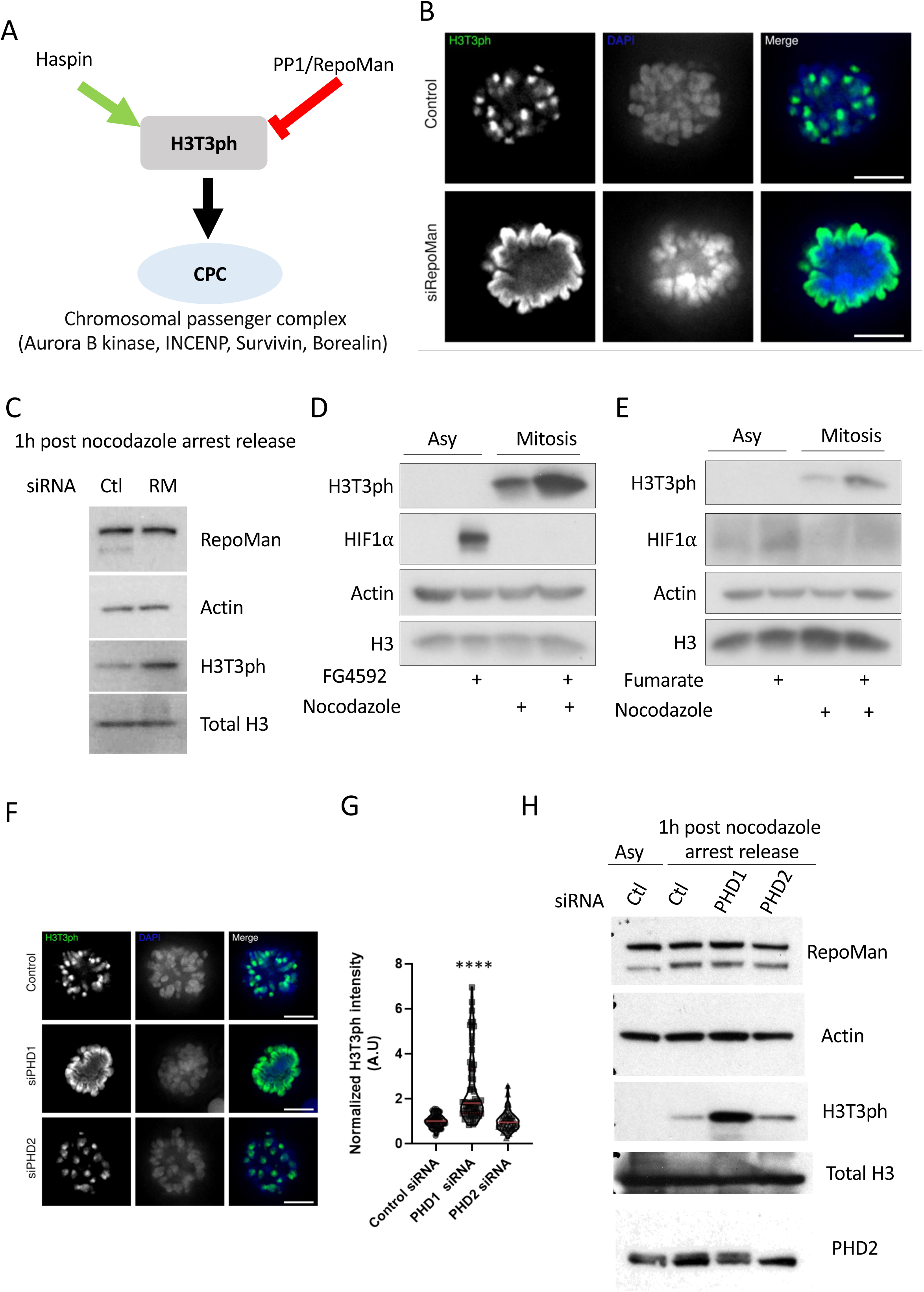
PHD1 regulates phosphorylation of H3T3 during prometaphase. (A) Schematic model of the role of PP1-RepoMan during prometaphase. (B) Hela cells were transfected with sicontrol , or siRNA RepoMan . Cells were arrested in prometaphase with nocodazole 100ng/ml for 16h and released from the arrest for 1h in normal media before fixation and stained with pH3T3 and DAPI. (C) Cells were treated as in (B) and harvested for western blot analysis (D) Asynchronous or prometaphase arrested HeLa cells were treated with FG4592 50 µM for 2h or DMSO. In case of mitotic cells, they were treated 1h before the release and during the release for another hour. Cells were lysed and subject to western blot (E) Asynchronous or prometaphase arrested HeLa cells were treated with 100µM Fumarate for 1h. In case of mitotic cells, they were treated for 1h during the release. Cells were lysed and subject to western blot. (F). Immunofluorescence images of HeLa cells arrested in prometaphase. HeLa cells were transfected with sicontrol, siPHD1, siPHD2 (as in B) Prometaphase arrested cells were release for 1h before fixation. Images scale bars represent 5µm. (G) Graph display the normalized pH3T3 intensity of 30 cells per condition of 3 independent experiments. pH3T3 intensity of each condition (siPHD1 and siPHD2) was normalized to the intensity values of pH3T3 in the sicontrol. The average of 3 independent experiments is shown. Unpair t test p<0.0001 Control vs siPHD1 (H) Western blot analysis of HeLa cells treated as in (F) and including asynchronous cells transfected with the control siRNA.

Next, we investigated the potential biological role of RepoMan P604 hydroxylation, analysing H3T3 phosphorylation during prometaphase. First, H3T3 phosphorylation was analysed after treating cells with the PHD inhibitor, FG4592. This revealed that FG4592 treatment resulted in increased levels of phosphorylated H3T3 in mitotic cells (Figure 3D), as does RepoMan depletion (Figure 3C). These observations are consistent with RepoMan hydroxylation being necessary for RepoMan-dependent dephosphorylation of H3T3 in cells during early mitosis (Figure 3D).

PHD enzymes are known to sense changes in levels of metabolites, as well as oxygen, for example, responding to increased levels of TCA metabolites, such as fumarate ^30^. Therefore, we investigated if PHD inhibition by fumarate also impaired the ability of RepoMan to regulate H3T3 phosphorylation. Treatment of HeLa cells with fumarate resulted in increased phosphorylation of H3T3 in mitotic cells (Figure 3E), analogous to that observed after FG4592 treatment. These data suggest that PHD activity is involved in the regulation of H3T3 phosphorylation during prometaphase and further indicate that prometaphase can be modulated in response to changes in levels of metabolites.

While we demonstrate here, and in the accompanying paper by Jiang et al ^18^., that PHD1 can hydroxylate RepoMan at P604, we wanted to investigate whether RepoMan may also be a target for PHD2. Therefore, we analysed levels of H3T3 phosphorylation by immunofluorescence, comparing how this was affected by siRNA knock-down of either PHD1, or PHD2. This showed that only depletion of PHD1, but not PHD2, resulted in increased levels of H3T3 phosphorylation in nocodazole-arrested cells, similar to that seen after siRNA-knockdown of RepoMan (Figure 3F-G). Similar results were observed when lysates of nocodazole-arrested cells, which had been depleted of PHD1 by siRNA, were analysed by immunoblotting (Figure 3H). Due to the lack of antibodies that can detect endogenous PHD1, we confirmed the knockdown of PHD1 in HeLa cells, either by immunofluorescence analysis of cells transiently expressing EGFP-PHD1, or by western blot analysis of a HEK293 cell line stably expressing GFP-PHD1 (Figure S1A-B).

Taken together, these results indicate that PHD1 is involved in RepoMan regulation and can modulate the levels of phosphorylated H3T3. The data are also consistent with the hydroxylation of RepoMan at P604 being important for its function during early mitosis.

### P604 hydroxylation is necessary for RepoMan function in early mitosis

To investigate further the functional role of RepoMan Pro604 hydroxylation in early mitosis, we generated stable HeLa cell lines that exogenously express either a YFP-tagged wt RepoMan or a YFP-tagged point mutation of RepoMan, in which the hydroxylated Proline 604 is replaced with Alanine, (YFP-RepoMan-P604A). These cell lines are doxycycline inducible and the YFP-RepoMan expressed is also resistant to siRNAs that deplete endogenous RepoMan, allowing for rescue experiments to be performed. Both the inducible RepoMan wt and P604A mutant expressed at comparable levels, similar to the levels of endogenous RepoMan (Figure 4A).

**Figure 4.**
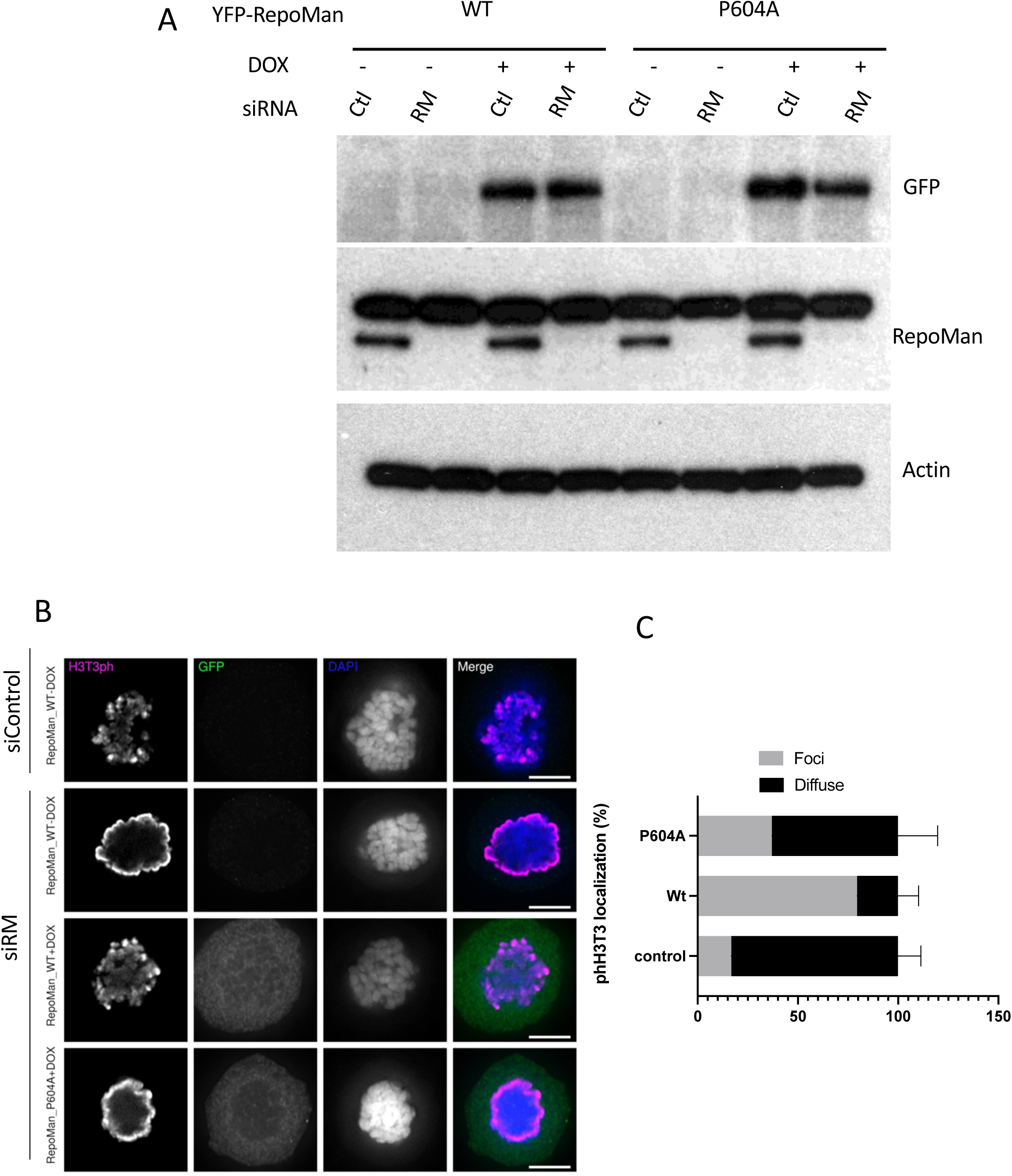
P604 in RepoMan is required for correct localisation and function in mitosis. (A) Western blot analysis of HeLa-YFP-RepoMan wt or the P604A mutant induction in presence of 1 µg/ml doxycycline. Cells were transfected with the siRNA of RepoMan and after 24h cells were induced with doxycycline for another 24h before harvested. Blots were developed with GFP, RepoMan and actin antibodies. (B) Immunofluorescence of HeLa-YFP-RepoMan wt or the P604A synchronized in prometaphase with nocodazole after siRM knockdown. Dox was used to induce the expression of the YFP proteins. Anti phH3T3, GFP antibodies were used, and DNA was stain with DAPI. (C) Quantification of (B) in cells treated with siRM and induced or not with doxycycline. The graph displays the distribution of phH3T3 on prometaphase cells. Graph represents the percentage of cells showing phH3T3 localization as foci (centromeric) or diffuse localization (along chromosomes arms). Average of 4 independent experiments with a total number of cells 46 for siRM (not induced), 46 for siRM + YFPRMwt and 52 for siRM + YFP-P604A (9-20 cells per condition per experiment).

We analysed by immunofluorescence the localization and levels of phosphorylated H3T3 to determine if this is affected during early mitosis by expression of the RepoMan P604A mutant protein. As expected, in control cells transfected with an siRNA control (without doxycycline), phosphorylated H3T3 localises preferentially to centromeres, while cells transfected with siRNA that depletes endogenous RepoMan show phosphorylated H3T3 spread along chromosome arms (Figure 4B-C; Figure S2).

The induction of YFP-RepoMan-wt in cells depleted of endogenous RepoMan by siRNA transfection clearly rescued the phosphorylated H3T3 phenotype observed in control cells, i.e., immunofluorescence again shows H3T3 localised preferentially to centromeres. Interestingly, however, induction with doxycycline of the YFP-RepoMan-P604A mutant, in contrast to the YFP-RepoMan-wt, was unable to rescue knockdown of endogenous RepoMan, with H3T3 phosphorylation still showing a similar distribution spread along chromosome arms as seen in cells depleted of endogenous RepoMan (Figure 4B-C; Figure S2). We note that this mislocalisation of phosphorylated H3T3, as seen both after siRNA depletion of endogenous RepoMan and in the presence of the exogenous YFP-RepoMan-P604A mutant, is similar to that observed in cells expressing endogenous RepoMan after siRNA mediated depletion of PHD1 (Figure 3F). Together, these results suggest that hydroxylation of RepoMan at P604 is required for the PP1-RepoMan complex to dephosphorylate H3T3 from chromosome arms during prometaphase.

### P604-RepoMan is required for PP2-B56 interaction during (pro)metaphase

The loading of PP1-RepoMan onto chromatin during mitosis is dynamic and depends on phosphorylation of Ser893 in the histone binding domain of RepoMan (Qian, 2015). This residue is phosphorylated by Aurora B and dephosphorylated by a pool of PP2A-B56 that is associated to RepoMan (Qian, 2015). As we saw no evidence that lack of hydroxylation on P604 alters levels of RepoMan protein (Figure S3A), we sought to determine whether the YFP-P604A RepoMan mutant mislocalized during the cell cycle, as compared with the equivalent YFP-RepoMan wt protein. To do this, we used live cell imaging in HeLa cells released from a 24h thymidine block (Figure S3B), comparing the degree of loading onto chromatin in early mitosis of the respective, exogenously expressed YFP-tagged wt and P604A RepoMan proteins. Both the wt and mutant constructs showed similar levels of loading onto chromatin during (pro)metaphase (Figure S3C).

We next compared the association of the YFP-tagged wt and P604A RepoMan proteins with chromatin as cells traversed mitosis by measuring the ratio of RepoMan to DNA signals in cells entering mitosis. We note that the level of RepoMan expression differs between individual cells analysed in this experiment, due to measurements being made with a mixed population of cells, rather than a clone. To account for this, we normalized the values of fluorescence intensity measured throughout mitosis progression, with the intensity of YFP-tagged RepoMan in G2 (T0), for each cell that was measured (Figure S3D). This analysis revealed that the YFP-RepoMan-P604A mutant loads onto chromatin almost twice faster than wt, suggesting potential changes to the interaction of RepoMan with B56/PP2A during prometaphase.

Phosphorylation of Ser893 on RepoMan, by AuroraB, prevents the loading of RepoMan to chromatin during prometaphase ^23^, with this phosphorylation being removed by the phosphatase PP2A/B56γ ^23,24^. The PP2A/B56 γ binding site on RepoMan occurs in the short linear motif (SLIM) ^31^ that is located 13 amino acids away from Pro604 (Figure 5A). We therefore hypothesised that the hydroxylation of Pro604 on RepoMan could affect the interaction with B56γ during prometaphase and thus promote the loading of the PP1-RepoMan complex to chromatin, leading to the dephosphorylation of H3T3 on chromosome arms. It has been reported that the interaction between B56γ and RepoMan takes place during mitosis, reaching a maximum at prometaphase ^24^.

**Figure 5:**
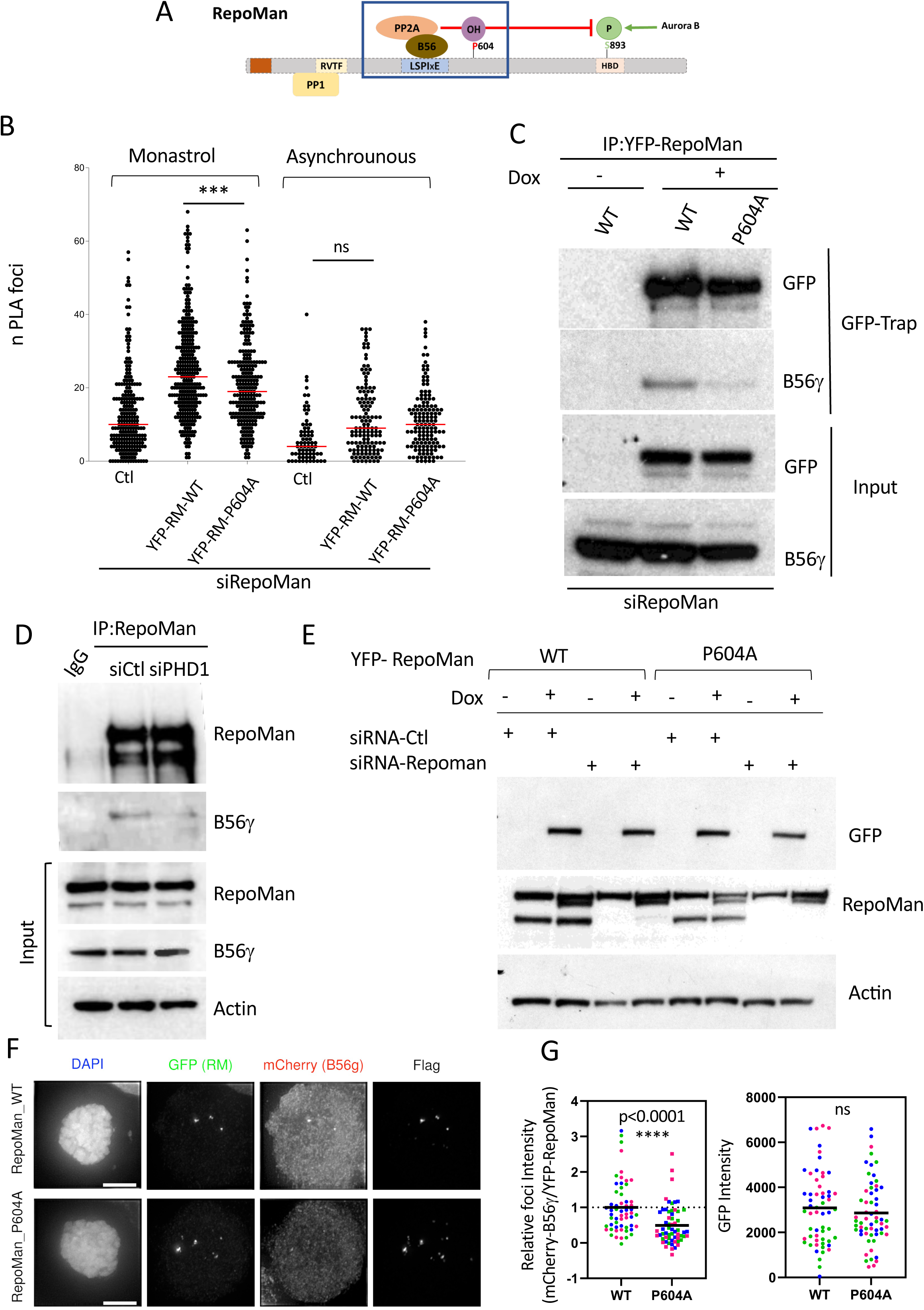
RepoMan P604 hydroxylation is required for the recruitment of B56γ in prometaphase cells. (A) Schematic representation of RepoMan Proline 604 proximity to B56-PP2A binding site (LSPIxE). (B) Proximity ligation assay. Graph represents the number of PLA foci per cell in the indicated conditions (monastrol arrested or asynchronous). Quantified PLA foci are located in the YFP area (RepoMan localisation) of the mitotic cells (phSer10 positive cells) Endogenous RepoMan was knocked down using siRM. Median is shown in red. Statistical analysis was performed applying an unpaired t-test and the p value comparing wt vs mutant upon monastrol treatment is <0.0001. Number of cells per condition in the same order than is shown are: 229,301,237,71,130 and 139. Before PLA reaction cells were fixed with PFA 4% and stained with Anti-GFP + B56γ antibodies or only GFP antibody was used as a negative control. After PLA reaction cells were stain with anti phH3Ser10 as a mitotic marker (C) Co-immunoprecipitation between YFP-RepoMan wt or P604A mutant with endogenous B56γ. Endogenous RepoMan was knocked down using siRM. 16h after transfection YFP RepoMan was induced with 1 µg/ml doxycline and cells were synchronized in prometaphase. Cells lysates were subject to immunoprecipitation using GFP trap magnetic beads and analysing by western blot. (D) Co-immunoprecipitation between endogenous RepoMan and endogenous B56γ in cells depleted or not of PHD1 using siRNA. Hela cells were transfected with siRNA (PHD1 or ctl), 24 h later cells were incubated with thymidine 2mM for another 24h. After the incubation time cells were release from the thymidine block, for 2h in normal media and arrested in prometaphase with nocodazole for 14h. Cells were harvested by shake off and lysed for immunoprecipitation of endogenous RepoMan. (E) Western blot analysis of HeLa-YFP-RepoMan wt or P604A / mCherry B56γ cell line. Cells were depleted of endogenous RepoMan and the indicated YFP-RepoMan variants were induced with dox. GFP, CDCA2 (RepoMan) and actin as a loading control were utilized. (F) Immunofluorescence showing the recruitment of mCherry B56γ through YFP-RepoMan wt or P604A at ectopic foci on chr 1 in prometaphase arrested cells. GFP, m-cherry and Flag antibodies were used to detect YFP, B56γ and dCas9 respectively. DAPI show DNA, Graphs represent B56γ levels (G) and GFP levels(G). Average of 3 independent experiments, 58 (wt) 60 (P604A) cells. Scale bar 5µm. Mann-Whitney test was applied P value <0.0001.

To test whether the hydroxylation of P604 on RepoMan alters its interaction with PP2A-B56γ during prometaphase, we performed PLA assays, (Proximity Ligation Reaction), in either asynchronous HeLa cell cultures, or in cells synchronised in prometaphase with monastrol (an inhibitor of Eg5), using anti-GFP and anti-B56γ antibodies to detect YFP-RepoMan and endogenous PP2A-B56γ, respectively. As shown by DAPI content, (n=G1 and 2n=G2/Mitosis), the cell cycle synchronization procedure was successful for both YFP-RepoMan-wt and the YFP-RepoMan-P604A mutant (Figure S4).

In prometaphase cells, YFP-RepoMan-wt shows a higher number of PLA foci, as compared with cells expressing the YFP-RepoMan-P604A mutant (Figure 5B). This is consistent with the YFP-RepoMan-P604A mutant having a reduced interaction with B56γ. To further validate this finding, co-immunoprecipitation experiments were performed, using GFP-Trap beads and cells synchronised with nocodazole in prometaphase, comparing expression of YFP-RepoMan-wt and the YFP-RepoMan-P604A mutant. While cells expressing the wt and mutant RepoMan constructs synchronised equally well (Figure S4), the YFP-RepoMan-P604A mutant showed a reduced interaction with endogenous B56γ, as compared with the YFP-RepoMan-wt protein (Figure 5C). In addition, in cells depleted of PHD1, endogenous RepoMan had a reduced interaction with endogenous B56γ (Figure 5D). Similar results were also observed with EGFP-RepoMan and endogenous B56γ (Figure S5).

We validated this important finding further, using an orthogonal approach. For this, cells were generated that expressed either YFP-RepoMan-wt, or the YFP-RepoMan-P604A mutant, (both constructs being doxycycline inducible and resistant to siRNA targeted to endogenous RepoMan), in cells that also constitutively express mCherry-B56γ. Western blot analysis of cell lysates transfected with either siRNA non-targeting control, or siRNA targeted to knock-down endogenous RepoMan, demonstrates that the expression levels of both exogenous YFP-RepoMan-wt and YFP-RepoMan-P604A proteins are comparable and similar to the level of endogenous RepoMan in these cells prior to knock-down (Figure 5E).

Next, we compared the interaction between YFP-RepoMan-wt and YFP-RepoMan-P604A, with mCherry-B56γ-wt on the telomere of chromosome 1, using the dead Cas9 (dCas9)-DARPin system ^32^. Cells were transiently co-transfected with dCas9-DARPin-Flag and sgChr1 expression vectors, following the depletion of endogenous RepoMan with siRNA. Both dCAS9-DARPin and the YFP-RepoMan constructs are doxycycline inducible. Cells were arrested in prometaphase with nocodazole and fixed for immunofluorescence analysis (see Methods). The GFP panel represents YFP-RepoMan, while the flag panel represents dCas-9-DARPin (Figure 5F). Upon recruitment of mCherry-B56γ, strong foci are seen when YFP-RepoMan-wt is induced (Figure 5F, upper row). However, in cells where YFP-RepoMan-P604A mutant is induced, the intensity of the foci is clearly reduced (Figure 5F, lower row). Quantification of the foci intensity (i.e., the B56γ/YFP ratio), confirms that there is a lower level of interaction between YFP-RepoMan-P604A and mCherry-B56γ, in comparison with YFP-RepoMan-wt (Figure 5G). Importantly, the YFP levels in foci are similar for expression of both the YFP-RepoMan-wt and YFP-RepoMan-P604A mutant, indicating that the difference observed in interaction with mCherry-B56γ is not due to differences in the level of YFP recruited to DARPin (Figure 5G).

Overall, these results indicate that Proline 604 in RepoMan is important for the interaction between B56γ and RepoMan during prometaphase and support a model in which the hydroxylation of RepoMan P604 favours interaction with B56γ in cells during prometaphase.

### Loss of RepoMan P604 Hydroxylation results in mitotic defects

Dephosphorylation of H3T3 by RepoMan-PP1 is essential for correct chromosome alignment and segregation ^28,33^. Since we show above that hydroxylation of RepoMan-P604 by PHD1 is important for efficient dephosphorylation of H3T3 during prometaphase, we analysed if chromosome alignment is affected by hydroxylation of RepoMan P604A. To address this, we performed a release from monastrol arrest (see Methods), which analyses correct establishment of spindle bipolarity and chromosome alignment ^34^. First, HeLa cells were arrested in prometaphase with monastrol (Figure S6), then released in fresh media and fixed at different time points after release and analysed by immunofluorescence (Figure 6A). Cells expressing either YFP tagged RepoMan-wt, or RepoMan-P604A, can form bipolar spindles after monastrol release (Figure 6B). However, the numbers of unaligned chromosomes observed are higher in bipolar cells expressing the YFP-RepoMan-P604A mutant, 1h after release from monastrol arrest (Figure 6C). Expression of YFP-RepoMan-P604A resulted in nearly 75% of cells displaying mitotic defects, a significant level compared to cells expressing YFP-RepoMan-wt.

**Figure 6:**
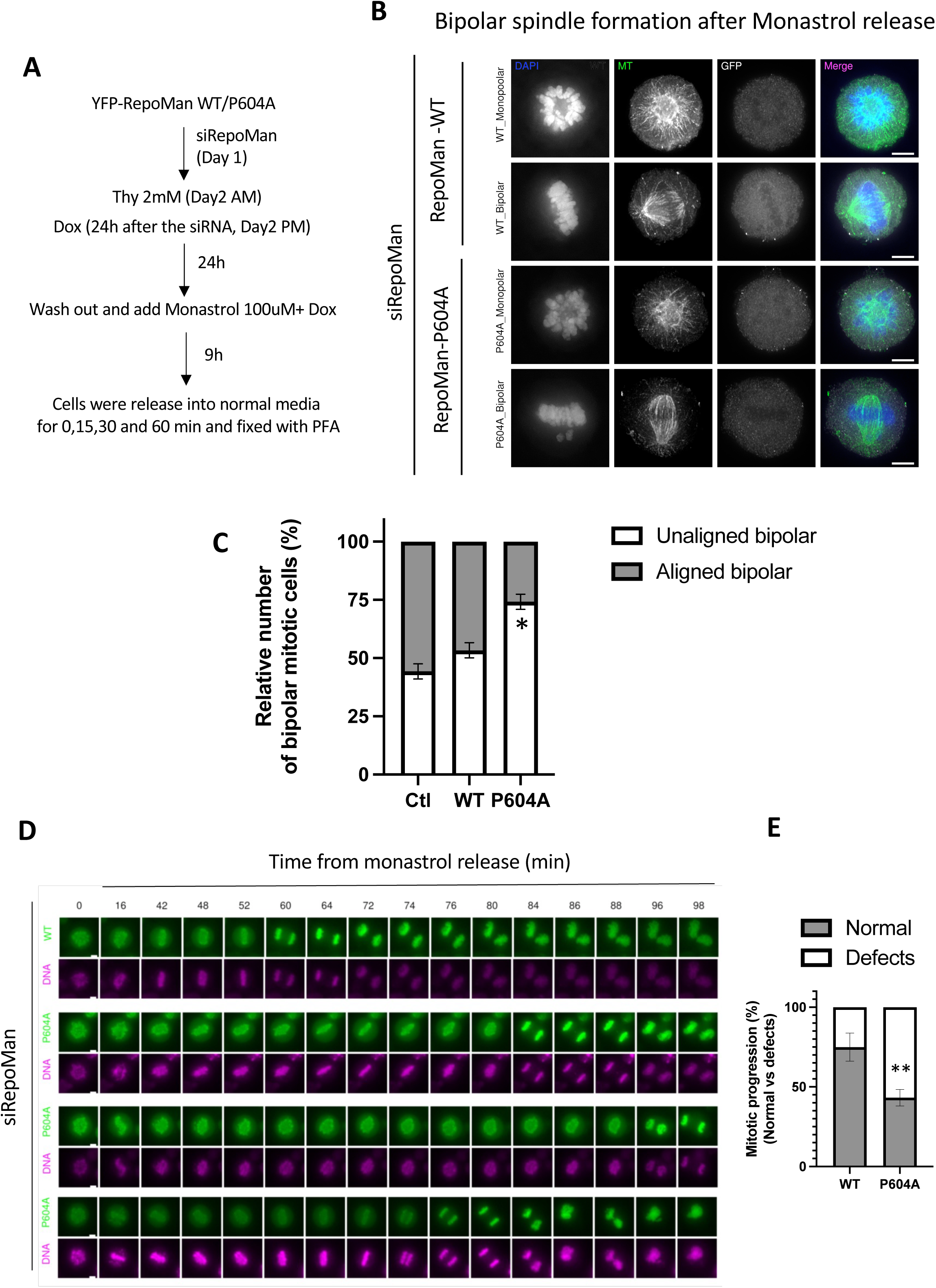
P604 of RepoMan is required for normal mitotic progression. (A) Schematic representation of the experimental design. (B) Representative Immunofluorescence of YFP-RepoMan arrested in prometaphase with monastrol. Cells were released for 30 min. Monopolar or bipolar cells are shown for either RepoMan wt or P604A. Cells were fixed and stained with MT, GFP antibodies and DAPI shows the DNA. Scale bar 5µm. (C) Quantification of chromosome alignment in bipolar spindles after release of 1h from monastrol arrest. The graph represents the percentage of cells with aligned or unaligned chromosomes in bipolar spindles respect to the total number of bipolar mitotic cells from 3 independent experiments is shown. Total cells analysed Ctl: 161, RM-WT 196, RM P604A 187. Unpaired t test unaligned wt vs mut p= 0.0105 (D) Time lapse of YFP-RepoMan wt or P604A cells arrested in prometaphase with monastrol and released into fresh media. (YFP-RepoMan) green and DNA (Magenta). Representative images are shown scale bar 5µm. (E) The graph shows the percentage of normal vs defective mitosis over the of total mitotic cells per condition per experiment. (Total number of cells analysed: 57 cells RepoMan wt and 55 cells for RepoMan P604A from three independent experiments). Unpaired t test p=0.0058 defects wt vs mut.

Using the same approach, we analysed mitosis progression in live cells. Time lapse analysis was performed in HeLa cells that had been arrested with monastrol in prometaphase (T=0) and then released into fresh media. On average, anaphase onset in cells expressing YFP-Repoman-wt occurred 60-70 mins after release from monastrol. However, in cells expressing the YFP-RepoMan-P604A mutant there was an increase in cells failing to complete mitosis, while most of those cells that completed mitosis after monastrol release did so more slowly, showing on average a delay of 15-30 mins compared with cells expressing YFP-RepoMan-wt (Figure 6D). Moreover, in cells expressing the YFP-RepoMan-P604A mutant, a variety of defects were apparent. For example, > 55% of cells displayed chromosome alignment and segregation problems, including an increased level of cell death (Figure 6D-E).

Taken together, these results support a model in which the hydroxylation of RepoMan at P604 is required for RepoMan function to ensure efficient progression through mitosis.

## Discussion

In the accompanying study by Jiang et al. ^18^, we have identified by mass spectrometry that the PP1 regulatory subunit, RepoMan (CDCA2), is hydroxylated at P604 by PHD1, both in cell lines and *in vitro*. Here, we have used a combination of cellular, biochemical and fluorescence imaging assays to analyse the potential functional significance of site-specific hydroxylation of RepoMan at P604. We have compared the effect of expressing wt RepoMan with expressing a single site mutation that replaces proline 604 with alanine. In addition, we have compared the effect of expressing the RepoMan-P604A mutant in cells, with siRNA knock-down of PHD1 in cells expressing wt RepoMan. Collectively, these experiments showed that replacing wild type RepoMan in cells with a RepoMan-P604A mutant resulted in major defects in early mitosis. These defects included mislocalisation of phosphorylated H3T3, along with decreased interaction with the B56 protein, which is an important targeting subunit for the PP2A-B56 phosphatase complex that dephosphorylates the PP1-RepoMan phosphatase to promote its loading onto chromatin, prior to anaphase onset. Furthermore, in cells expressing the RepoMan-P604A mutant a variety of defects were apparent, including problems with chromosome alignment and segregation and increased cell death, as previously seen in cells expressing endogenous RepoMan after depletion of PHD1^19^

In combination, the data summarised above lead us to propose the model shown in Figure 7., In this model, the PP1-RepoMan phosphatase complex associates with chromatin throughout the cell cycle, except during prophase when cells enter mitosis. During prometaphase, PHD1-mediated, site-specific hydroxylation of RepoMan at P604 (OH-P604), contributes to stabilising the interaction between the PP1-RepoMan and PP2A-B56 phosphatase complexes, thereby allowing PP2A-B56 to dephosphorylate RepoMan at S893, which in turn allows re-loading of PP1-RepoMan onto chromatin. We propose that when the PP1-RepoMan complex is bound to chromatin, it dephosphorylates phH3T3 on chromosome arms, which aids in targeting the chromosomal passenger complex (CPC) to preferentially bind to centromeres, where higher levels of phH3T3 remain and act as docking sites for the CPC. The PP1-RepoMan phosphatase complex then remains associated with chromatin throughout the remainder of mitosis and the following interphase. Entry into the next mitotic cycle sees the activation of Aurora B kinase, which phosphorylates RepoMan on Ser893 ^23^ and prevents its association with chromatin during prometaphase.

**Figure 7:**
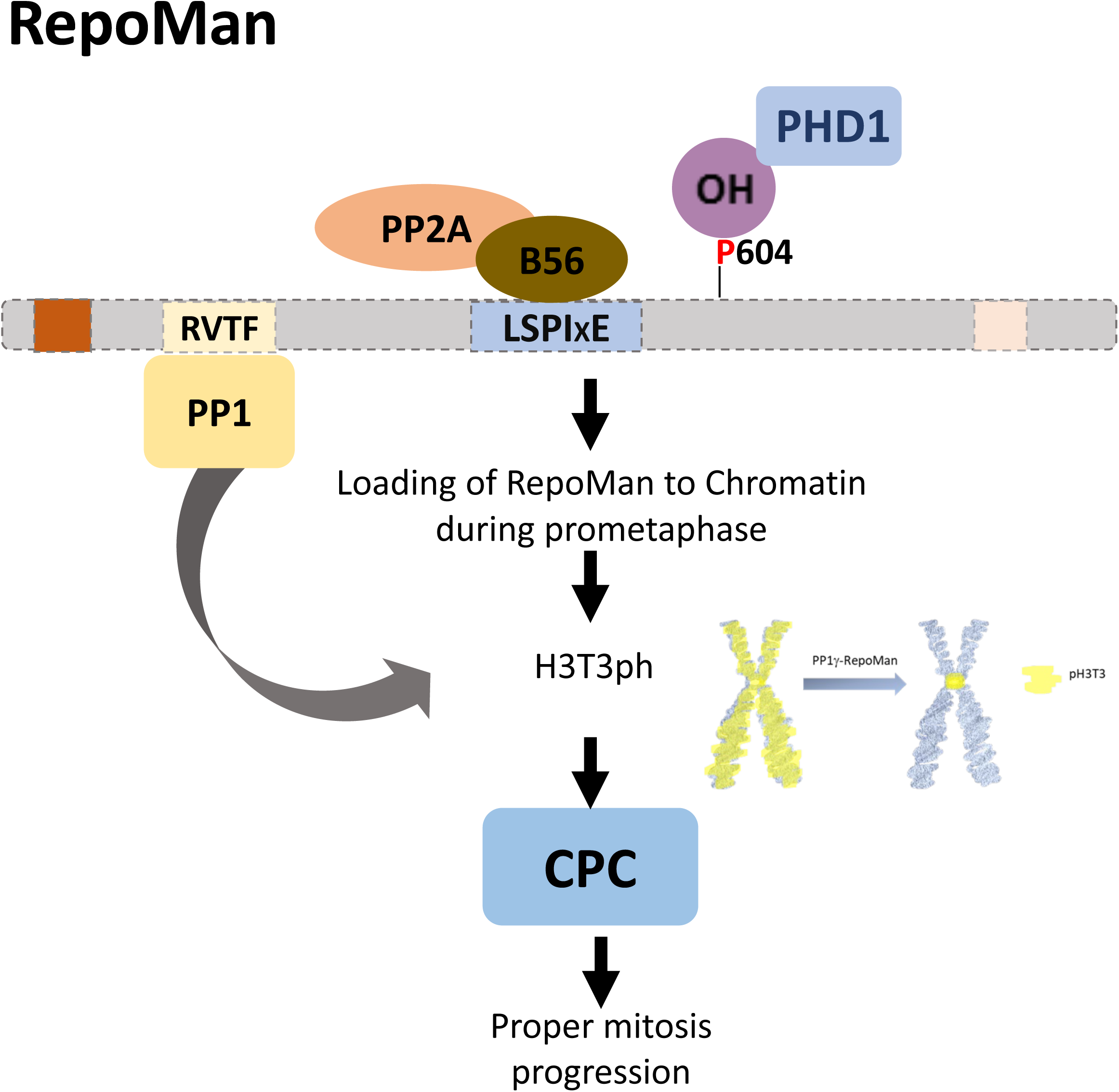
Schematic model of RepoMan hydroxylation during prometaphase. PHD1 hydroxylates RepoMan at proline 604. During prometaphase this modification is important for the binding of PP2A-B56γ to RepoMan. PP2A-B56γ has a crucial role in the loading of RepoMan to chromatin during prometaphase, leading PP1-RepoMan to dephosphorylates phH3T3 from chromosome arms. The resulted enrichment on the phH3T3 at the centromere assures the correct localisation of the CPC (chromosomal passenger complex) important for the proper mitosis progression.

To the best of our knowledge, the model shown in Figure 7 is consistent with all experimental observations, both from our current work and from previous studies by ourselves and others, concerning the dynamic localisation of the RepoMan protein during the cell cycle and its role in mechanisms controlling progression through mitosis ^21,22,24,33,35–37^. Importantly, the data here and in the accompanying manuscript by Jiang et al. ^18^, use mass spectrometry to identify unambiguously the site-specific hydroxylation of RepoMan at P604 by PHD1 and then characterise the functional significance of this post translational modification for protein interactions involved in mechanisms important for efficient control of mitotic progression. These new data support and extend our previous observations that PHD enzymes contribute to the regulation of cell cycle progression ^17,19,20^. Furthermore, these data identify specific molecular mechanisms whereby both oxygen sensing mechanisms and the concentration of specific metabolites, such as fumarate, can influence the cell cycle, by affecting the activity of PHD enzymes and changing their ability to hydroxylate proline residues on novel target proteins whose modification state affects their functions in cell cycle control. This provides a new paradigm for understanding mechanisms that can integrate cell cycle regulation in cells and tissues with signals reporting on exposure of organisms to stress and other physiological cues important for homeostasis.

Interestingly, PHD-dependent proline modification can also be modulated in concert with other post translational protein signalling mechanisms, including phosphorylation ^16^. Of note, P604 is next to T603, which has been shown to be phosphorylated in mitosis in previous phosphoproteomic screens ^38,39^. It is therefore possible this phosphorylation might be impacted by P604 hydroxylation. However, additional work is needed to investigate this further.

One interesting aspect of this work is the specificity for PHD1 over PHD2. For HIF regulation, PHD2 is dominant, with PHD1 and PHD3 having lesser roles ^7^. As our knowledge of PHD-targeting and specificity is still based on HIF, more analysis is necessary to really understand how individual PHDs are controlled and select their targets. One possibility is direct modification of PHD enzymes themselves by for example phosphorylation. Phosphorylation of PHD1 by CDKs could alter target selectivity between HIF and Cep192 for example ^20^, however more work is needed to fully understand the regulation of these enzymes in cells.

We note that in addition to proteins involved in cell cycle regulation, our MS analyses of PHD target proteins has also identified sites of proline hydroxylation in many new proteins that have roles in other important cellular processes, notably including RNA processing ^18^. It will be interesting in future, therefore, to pursue the functional characterisation of these novel PHD targets, to help deepen an understanding of the breadth of cellular mechanisms in which post-translational modification of proteins via proline hydroxylation can modulate function.

There has been a degree of controversy regarding the physiological relevance of new PHD targets identified in cells, beyond HIFα, due, at least in part, to the reported lack of activity of PHDs *in vitro* towards hydroxylation of non-HIF1 peptides ^40^. Nonetheless, evidence has been growing from a multitude of laboratories in support of functional roles for site-specific proline hydroxylation by PHDs on protein targets other than HIF1α ^1,16^. For example, Histone H3 ^25^ and Beclin ^41^ for PHD1, AMPK ^42^ and IRF3 ^43^ for PHD2. The data in this study clearly show the functional importance of PHD1-dependent hydroxylation of RepoMan for efficient mitotic progression, providing an important example that PHD targets are not confined to only HIF1α.

## Materials and methods

### Cell Culture

HEK293, HeLa ,HeLa Flp-in cells ,HeLa YFP-RepoMan WT siRNA Resistant ^44^ and HeLa mCherry-B56γ ^32^, were cultured in Dulbecco’s Modified Eagle Medium (Gibco, # 41966-029) supplemented with 10% Fetal bovine Serum (FBS,Gibco # A3169801), 100 U/mL penicillin and streptomycin (PS) and 2mM L-Glutamine. Cell lines were maintained at 37°C with 5% CO_2_ in a humidified incubator. During fluorescence time-lapse analysis cells were cultured in Leibosvitzs L15 media (Gibco #21083-027) supplemented with FBS and PS. HeLa Flp-in and HeLa mCherry-B56γ cells were used to stably express doxycycline inducible constructs after transfection with the relevant pcDNA5/FRT/TO vector and the Flp recombinase pOG44 (Invitrogen). Cells were then selected for stable integrants at the FRT locus using 200µg/mL hygromycin B (Roche) for 2 weeks. A pool of cells generated (rather than a single clone) was used in the experiments in this study.

### Cell cycle synchronisation

Unless indicated otherwise, a prometaphase arrest was induced by culturing cells consecutively for 24h with 2mM thymidine, 2h without thymidine and either 16 h with 100ng/mL nocodazole, or 6-9h with 100µM monastrol. The arrested cells were harvested by shake-off for biochemical analysis. Doxycycline (1µg/mL; Sigma-Aldrich), was used to induce expression of the YFP-RepoMan constructs, during and after the thymidine block, after 16-24h of siRNA knock-down to deplete endogenous RepoMan.

### Reagents

Final concentrations used: Doxycycline 1µg/mL, Nocodazole 100ng/ml (Sigma), Monastrol 100µM (Torsis), Thymidine 2mM (Sigma), Roxadustat (FG-4592, Selleck # S1007), Hygromycin B (200µg/µL) (Roche), Sir DNA far-red (1:10000) from Spirochrome SC007. Poly-L-lysine solution (P4707, Sigma) was used to coated coverslips for immunofluorescence.

### Plasmids and Mutagenesis

pEGFP-N1_PHD1 (Addgene #21400). dCas9-DARPin-Flag and pU6-sgChr1 ^32^, pcDNA5-YFP-RepoManWT expressing an siRNA-resistant and N-terminally YFP-tagged WT RepoMan ^44^. This vector was used to generate pcDNA5-YFP-RepoMan^P604A^ using the site directed mutagenesis kit (Q5#E0552S NEB), with the following primers: Forward 5’-TGAGATGACACGTTCCATTCCGAG-3’, Reverse 5’-GGGACTTCAGGCTCG-3’.

#### Interference RNA transfection

Double-stranded interference RNA (siRNA) was transfected at a final concentration of 20-30nM using the RNAi max Transfection Reagent (Invitrogen), according to the manufacturer’s instructions, in a 6 well plate. siControl, siPHD1 and siPHD2 sequences were described in (Moser et. al., 2013^19^). RepoMan (5’-UGACAGACUUGACCAGAAA-3’). All siRNAs were obtained from Eurofins genomics and synthesised with a dTdT overhang. For all experiments with either HeLa-YFP-RepoMan-wt, or the HeLa-YFP-RepoMan-P604A mutant, the endogenous RepoMan mRNA was first knocked down using siRNA, allowing replacement with the doxycycline-inducible, siRNA-resistant RepoMan constructs.

#### DNA transfection

HeLa cells were plated in either 6cm, or 10cm plates, 24h before transfection. Cells were transfected with either 1µg, or 5µg, of plasmid DNA EGFP-PHD1 using JetPRIME reagent (Ratio 1:2) (Polyplus), according to the manufacturer’s instructions, in media without antibiotic. 4h after transfection, the media was changed to complete media and after 48 hours further incubation cells were harvested in Lysis buffer. For siRNA treatment, HeLa cells were co-transfected in 6 well plates with the siRNA and 16h later with 1µg of plasmid DNA (EGFP-PHD1) for another 48h (Fig S1A-B). Cells were split to coverslips after siRNA and DNA was transfected. For Figure 5, DNA was transfected with FuGene (Promega), as described below.

#### Immunoprecipitation

Asynchronous, or prometaphase arrested HeLa cells were resuspended in 200µL of lysis buffer (50mM Tris pH 7.5, 150mM NaCl, 1% NP40, 0.5% Sodium deoxycholate and 0.1% SDS), containing protease inhibitors (Roche, Complete Mini EDTA-Free) and phosphatase inhibitors (PhosSTOP, Roche). Lysates were incubated on ice for 20 min and cleared by centrifugation at 4°C for 15 min at 13,000g. Supernatant was diluted in 300µl of Dilution buffer (50mM Tris pH 7.5, 150mM NaCl) 10% of material was taken as input. 500µL of the diluted lysate was transferred to a new tube containing pre-washed GFP-Trap magnetic agarose beads (ChromoTek) and incubated for 2h at 4°C. After the incubation time, beads were washed 3 times with a washing buffer (50mM Tris pH 7.5, 150mM NaCl and 0.05% NP40) and transferred to a clean tube after the last wash. Beads were resuspended in LDS sample buffer (Invitrogen), heated at 70°C for 10 min and the eluted proteins were transferred to a new tube. For immunoprecipitation of endogenous RepoMan, lysates were incubated with 1.5 µg of CDCA2 antibody (or IgG as a control) ON at 4C. Antibody-Antigen complex was incubated with magnetic Dynabeads A beads (Invitrogen) for 2h at 4C. Eluted proteins were analysed by Western blot.

### Western blot and dot blot

Western blot analysis was performed in NuPAGE 4-12% Bis-Tris gels (Invitrogen), or normal acrylamide gels. Immobilon-P transfer membranes (PVDF, 0.45µM pore size, Millipore) were used for a semi-dry transfer. After blocking in 10% fat-free milk in TBS-tween, membranes were incubated overnight with the primary antibody. After 3 washes with TBST plus one with TBS for 5 min each, membranes were incubated with the HRP secondary antibodies and developed using Pierce-ECL western blotting substrate (32106, Thermo Scientific).

### Antibodies

The following primary antibodies were used for either western blot, or IF, as indicated.

CDCA2 (HPA030049, Sigma) 1/1000 WB, 1/300 IF, Phospho Histone 3 (Thr3) (13576S, Cell signaling) 1/3000 IF, Phospho Histone 3 (Thr3) (07-424, Millipore) 1/1000 WB, Histone 3 (9715S, Cell signaling) 1/1000 WB, Actin (3700S, Cell signaling) 1/5000 WB, PHD2/Egln1 (4835S, Cell signalling) 1/1000 WB, HIF-1α (610959, BD Biosciences) 1/1000 WB, ,PP2A-B56γ (A11) (sc-374379, Santa Cruz Biotechnology) 1/1000 WB , 1/100 PLA, GFP rabbit IgG fraction (A11122, Invitrogen) 1/100 PLA, GFP (Ab13970, Abcam) 1/1000 IF, GFP (47859600, Roche) 1/1000 WB. mCherry (GTX 128508, GeneTex) 1/1000 IF, αTubulin, clone DM1A (T9026, Sigma) 1/500 IF, Flag M2 (F3165, Sigma) 1/2000 IF. Fluorescently labelled secondary antibodies for Immunofluorescence were obtained from Jackson immunoresearch (1/250) and Invitrogen (1/1000) A488 goat anti-chicken (A11039), A568 goat anti-rabbit (A11036) and A647 donkey anti-mouse (A31571). For Western blot HRP secondary antibodies were used at 1/5000 Anti-rabbit and anti-mouse IgG-HRP linked (7074S and 7076S cell signalling) respectively.

### Proximity ligation assay (PLA)

Cells were transfected in 6 well plates with siRepoMan using Lipofectamine RNAimax (#13778-150 invitrogen). 14h later cells were split in media containing 2mM thymidine and 1µg/ml Doxycycline (to induce the expression of the YFP constructs) and transferred to a 96 well plate clear base (Greiner 655090) pre-coated with poly L-Lysine solution (sigma #P4707). After 24h with thymidine, cells were washed once with PBS and released in media containing 1µg/ml doxycycline and 100µM monastrol for 9h. Cells were fixed with 4% PFA for 10 min at room temperature and washed with PBS. Cells were incubated with 3% BSA in PBS plus 0.1% Triton for 30 min at room temperature and washed with PBS. From this point fixed cells were subject to Duolink PLA Fluorescent protocol according to the manufacturer’s instructions (Sigma), with some modifications. Briefly: cells were incubated with 40µL/well blocking solution 1x (DUO82007) for 45 min at 37⁰C in a humidified chamber and then incubated overnight at 4⁰C with the primary antibody (anti-GFP (1/100) plus anti-B56γ antibody diluted 1/100, or only with anti-GFP (diluted 1/100) as a negative control) in Duolink antibody diluent. After overnight incubation at 4°C in a humid chamber, samples were washed 2 times with buffer A at room temperature for 5 min. Anti mouse and anti-rabbit +/- probes were added 1:5 in Duolink antibody diluent and incubated for 1h at 37⁰C. After 2 washes with buffer A at RT the ligase was added for 30 min at 37⁰C, washed twice at RT with buffer A and polymerase together with the red fluorescent reagent was added for the amplification reaction for 100min at 37⁰C. After incubation, cells were washed at RT for 10 minutes with buffer B, 10 min with DAPI 1/1000 in PBS, 10 minutes with buffer B and 1 minute with 1/100 buffer B. After PLA assay, phSer10 antibody and secondary Alexa 647 were added to ensure that quantified cells were mitotic cells. We got similar results avoiding this step.

High throughput microscopy images were taken and analysed with ScanR High Content Screening Microscopy (Olympus). PLA foci per cell were detected with the foci detection package. Data shown in this work correspond with the PLA signal detected in the YFP area of each cell, that corresponds with the RepoMan signal. In addition, cells were filtered by phospho-Serine 10 positive signal. DAPI signal was used to classify cells according to their cell cycle stage (Figure S3). Data were visualised and statistically analysed in Tableau and GraphPrism. The statistical test was unpaired *t*-test.

### dCas-9-DAR protein-protein interaction assay

The interaction of either YFP-Repoman-WT, or YFP-Repoman-P604A mutant with mCherry-B56γ was investigated using a previously described interaction assay ^32^. Experiments were performed by transfecting YFP-RepoMan (wt or P604A) mCherry B56γ cells with dCas9-DARPin-Flag and pU6-sgChr1 (ratio 1:3) using Fugene (Promega) in a 6 well plate. 24h later cells were split and transfected with siRepoMan RNA before replating cells in 24 well plates on coverslips. 16h later cells were incubated with 2mM Thymidine plus 1 µg/ml doxycycline for 24h. After incubation cells were released from the thymidine block in media containing 3.3µM nocodazole and 1 µg/ml doxycycline for 8h and fixed with 4% PFA at RT for immunofluorescence. Cells were stained with antibodies to detect YFP-RepoMan, mCherry-B56γ and DARpin-Flag. Only mitotic cells that show a clear spot of YFP-RepoMan were imaged and quantified.

### Immunofluorescence

Cells on High Precision 1.5H 12 mm coverslips (Merienfield) were fixed with 4% PFA in PBS for 10 min at room temperature. Fixed cells were washed 3 times with PBS and incubated with 3% BSA in PBS + Triton 0.1% for 30 min at room temperature. Coverslips were incubated overnight at 4^⁰^C with primary antibody in PBS + 3% BSA. After 3 washes with PBS, coverslips were incubated for 2-3 h at room temperature in the dark with secondary antibody plus DAPI (D9542, Sigma 1/1000) in PBS + 3% BSA. After incubation, coverslips were washed three times with PBS for 10 min at RT. Each coverslip was mounted on slides using Prolong gold antifade reagent (Invitrogen #P36930). All images, apart from those shown in Figure S1A, were acquired on a Deltavision fluorescence microscope (Applied Precision, Inc, Issaquah, WA) using a 100x objective, 1x1 binning and processed using SoftWoRx software. Images shown in Figure S1A were similarly acquired on a Deltavision fluorescence microscope but using a 60x objective. Figure panels were created using OMERO ^45^.

### Time-lapse analysis

Cells were plated in iBIDi µslide 8 well plates and imaged in L-15 media, using a DeltaVision system equipped with a heated 37°C chamber, with a 40x objective, 4x4 binning, 256x256 image size, 4 z-stacks of 5µm each (total thickness 20µm), using softWoRx software (Applied Precision, Issaquah, WA). DNA was stained using SiRDNA a live cell far red fluorescent dye (Spirochrome, #SC007). Before adding the monastrol, after the thymidine block, cells were incubated with sirDNA 1/10,000 for 20 min, then the media was replaced with media containing Monastrol for the indicated time. Cells were released from the arrest and immediately imaged. For experiments shown in Figures S3B-C, after cells were released from thymidine, SirDNA was added for 20 min before imaging.

### Quantitative analysis of immunofluorescence and chromatin loading

For immunofluorescence data, the following macro was run in Fiji to measure background-corrected nuclear intensity in 2 channels. Individual nuclei were identified at user-selected points of interest by auto-thresholding using the Triangle algorithm <<REF-GB3>> followed by “Fill holes”. A user-selected region of interest (ROI) was used to estimate background and calculate background-corrected total intensity for both channels, as well as normalised intensity of the second channel relative to the nuclear stain. A second macro was used to prepare data for tracking after deconvolution using SoftWoRx: 1) images were cropped to remove border artefacts, 2) for each timepoint an average projection over in-focus Z slices was performed, and 3) background subtraction was performed using a sliding paraboloid with radius=50 pixels. A final macro was used to track nuclei semi-automatically and extract intensity data for loading analysis. Single Z-section time series were tracked starting from a user-defined rectangular selection containing a single nucleus of interest. At each timepoint, a nucleus selection was identified by Otsu auto-thresholding <<REF-GB4>>, fill holes and optional watershed to split touching objects (a cost function based on area and position changes determines whether the watershed operation is beneficial). Parameters used were: minimum and maximum nucleus area (45-500µm^2^) and maximum frame-to-frame displacement (20 pixels, 12.68µm) corresponding to ∼0.05µm/sec. The nucleus selection was used as the ROI to measure intensity statistics in a second channel corresponding to Repo-Man protein. Analysis of image data was carried out using an ImageJ<<REF-GB1>> macros in Fiji<<REF-GB2>>.

For quantification of GFP-PHD1 and RepoMan colocalization, Pearson correlation was calculated using the established method by Costes et al ^26^ and the JACoP plugin macro for Fiji ^46^. using Costes automatic threshold.

For quantification of images shown in Figure 5F an ImageJ macro was designed to threshold and select the Chrl foci automatically and to measure the mean foci intensity of mCherry -B56 relative to YFP-RepoMan, as previously described in ^32^.

## Authors contributions

JD designed and performed most of the experiments, analysed the data and wrote the draft of the manuscript; HJ conducted mass spectrometry experiments, analysed the data, and edited the manuscript; DS, FC and VA performed experiments, MP, AC analysed the data, CA and AS were involved in supervision and data analysis, JRS, SR, and AIL, designed the project, supervised, acquired the funding, analysed the data, drafted and edited the manuscript.

## Acknowledgements

We would like to thank members of the labs involved for helpful discussions and Dr. Graeme Ball (Dundee Imaging Facility) for help in analysing microscopy images. This work was supported mainly by the Wellcome Trust (JD, HJ, DS, FC, JRS, AIL and SR) (206293/Z/17/Z). AC was funded by a Wellcome Investigator grant to A.T.S. (222494/Z/21/Z). Research is CA lab is supported by the ERC-Stg-IDRE. VA is supported by the CRUK-CDF C57404/A21782. AIL acknowledges additional funding from BBSRC Ref: BB/V010948/1; Ref: APP3732 & UKRI Ref: EP/Y010655/1.

