## supplementary figures and legends for "PHD1-dependent hydroxylation of RepoMan (CDCA2) on P604 modulates the control of mitotic progression" (from another version of this preprint: version 3, 2025-11-30)

**Supplementary Figure Legends:**

**Figure S1: siPHD1 is effective at depleting PHD1 protein.**

(A) Immunofluorescence analysis of siPHD1 knockdown in HeLa cells transiently transfected with EGFP-PHD1. sicontrol was used as a control. GFP antibody was used for EGFP-PHD1 and DNA was stained with DAPI. (B) Western blot analysis of siPHD1 or si PHD1 UTR knockdown on HEK293 cells stable expressing GFP-PHD1 or GFP empty vector.

**Figure S2: siRepoMan is effective at depleting endogenous but not exogenous RepoMan.**

Hela cells were transfected with siRNA control or siRNA against endogenous RepoMan (siRM). 24h later cells were incubated with thymidine 2mM +/- doxycycline for 24 h. After the incubation time, cells were release from the thymidine for 2 h in normal media and nocodazole +/- dox was added for 14-16h. Cells were lysed and protein extracts analysed by western blot.

**Figure S3: Wt and P604A mutant RepoMan can load to chromatin.**

(A) Western blot analysis of endogenous RepoMan levels in presence of PHD inhibitor FG4592 50 μM for the indicated times. DMSO was used in time 0 as a control for 24h. MG132 20 μM was added for 3h. CDCA2, HIF1α and Actin antibodies were used. B) Cells were transfected with siRNA control or si RM. 16h later cells were incubated with thymidine 2mM +/- doxycycline to induce YFP-constructs or not (siCtl – dox) for 24 h. After the incubation time cells were release from the thymidine block for 6 h in normal media. Cells were lysed and analysed by western blot. (C) Both RepoMan wt and P604A mutant can load to chromatin during pro(methaphase). Live cell images of YFP RepoMan wt (Upper) or P604A mutant in metaphase. Cells were depleted of endogenous RepoMan with siRNAs., Cells were incubated with thymidine plus doxycycline for 24h and released in fresh media + doxycycline. Before imaging DNA was stained with sirDNA for 20 min in L15 media, after the incubation time, the media containing sirDNA was removed and cells were incubated in L15 media for cell imaging. (D) Graph depicting tracking of YFP-RepoMan wt or P604A mutant over chromatin during mitosis. Cells were treated as in (C). The graph represents the ratio between Repoman/DNA intensity over mitosis, values of 2 independent experiments (total number of cells between 2 experiments: RepoMan wt 29 and RepMan P604A 24. The values were normalized to the Repoman/DNA intensity during interphase 2 frames before cells enter in mitosis.

**Figure S4: Wt and P604A RepoMan cells are synchronised by monastrol.**

Cell cycle distribution of the experiment shown. The number of cells is represented in the y axis and DAPI content in the x axis, from n (G1) to 2n (G2/mitosis), marked with dash lines. Fist line of graph show asynchronous cells and the graph on the bottom shows monastrol treated cells. The whole cell population from each well was analysed. The number of cells per sample: Asynchronous 3668,7725,7621; monastrol 5089,5197,6820. One representative experiment is shown.

**Figure S5: PHD1 is required for the interaction between GFP-RepoMan and endogenous B56γ in mitosis.**

Hela cells were transfected with siRNA (PHD1 or ctl). Next day cells were transfected with 5µg of EGFP-RepoMan for 4 h. After the incubation time, cells were incubated with thymidine 2mM for 24h, release for 2h and incubated with nocodazole 100ng/ml for 14 h. Cells were harvested by shake off and resuspended in Lysis buffer. Lysates were subjected to GFP Trap IP and the interaction of overexpressed EGFP-RepoMan with endogenous B56γ was analysed.

**Figure S6. Efficient depletion of endogenous RepoMan and rescue with exogenous RepoMan wt and P604A.**

Hela cells were transfected with siRNA control or si RM. 16h later cells were incubated with thymidine 2mM +/- doxycycline to induce YFP-constructs or not (siCtl – dox) for 24 h. After the incubation time cells were release from the thymidine block for 9h in media containing Monastrol with or without doxycycline as indicated. After the incubation time cells were lysed and extracts analysed by western blot.

Figure S1

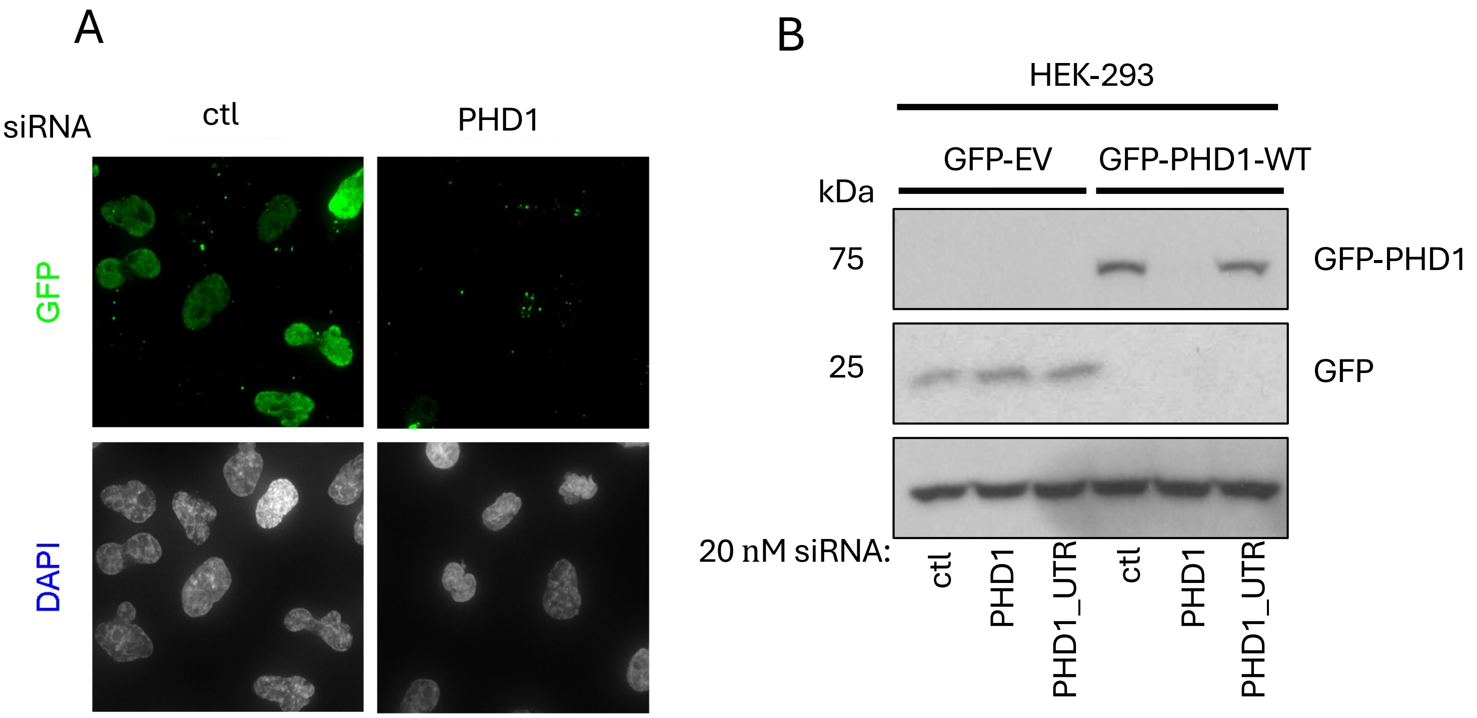
A
siRNA
ctl
PHD1
GFP
DAPI
B
HEK-293
GFP-EV
GFP-PHD1-WT
kDa
75
25
GFP-PHD1
GFP
20 nM siRNA:
ctl
PHD1
PHD1_UTR
ctl
PHD1
PHD1_UTR

Figure S2

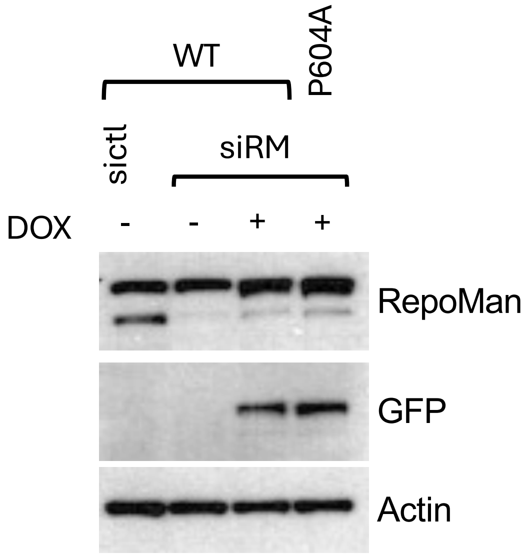

Figure S3

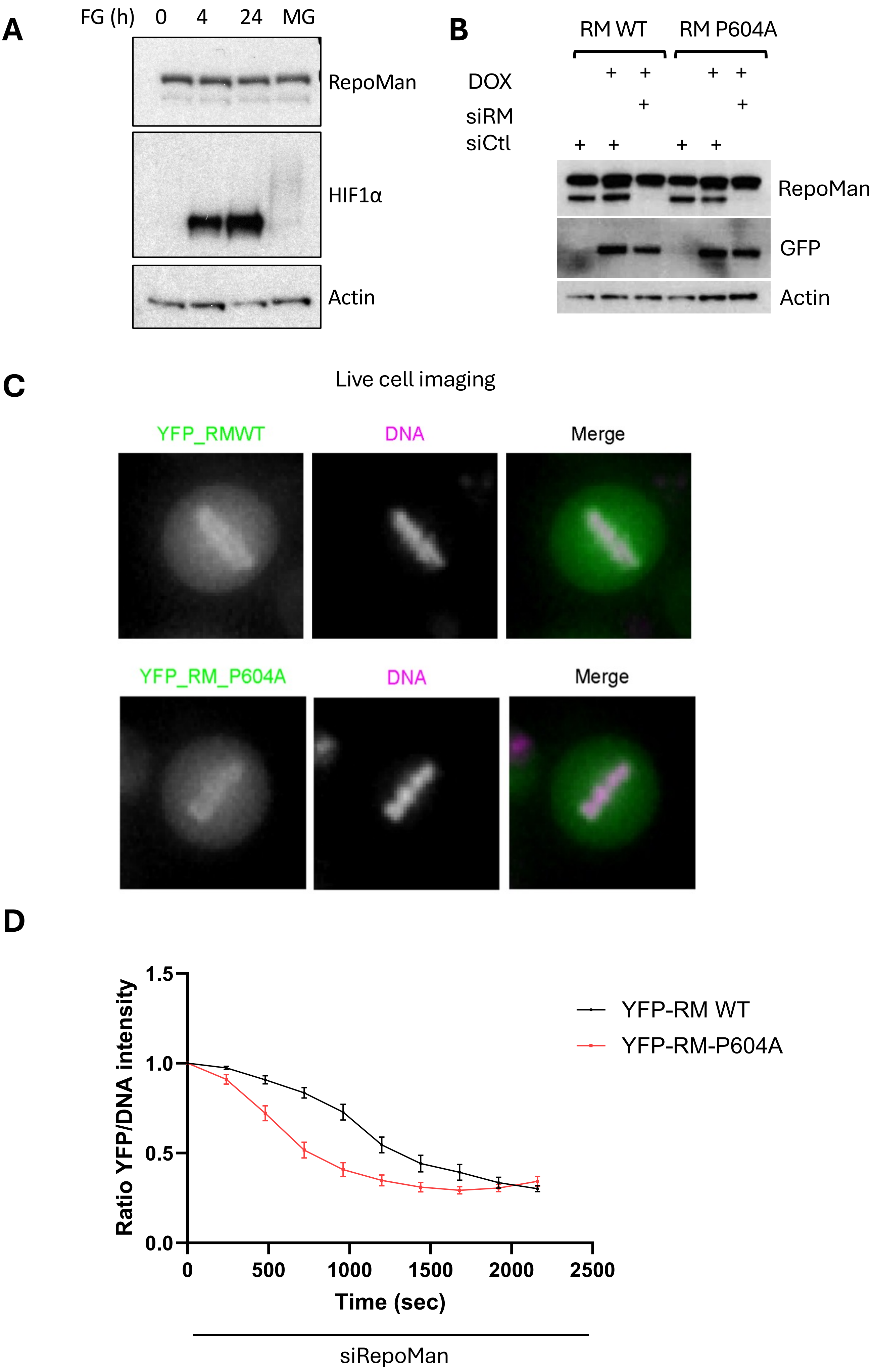
A
FG (h) 0 4 24 MG
RepoMan
HIF1α
Actin
B
RM WT
RM P604A
DOX
siRM
siCtl
RepoMan
GFP
Actin
C
Live cell imaging
YFP_RMWT
DNA
Merge
YFP_RM_P604A
DNA
Merge
D
Ratio YFP/DNA intensity
1.5
1.0
0.5
0.0
0
500
1000
1500
2000
2500
Time (sec)
YFP-RM WT
YFP-RM-P604A
siRepoMan

### Figure S4

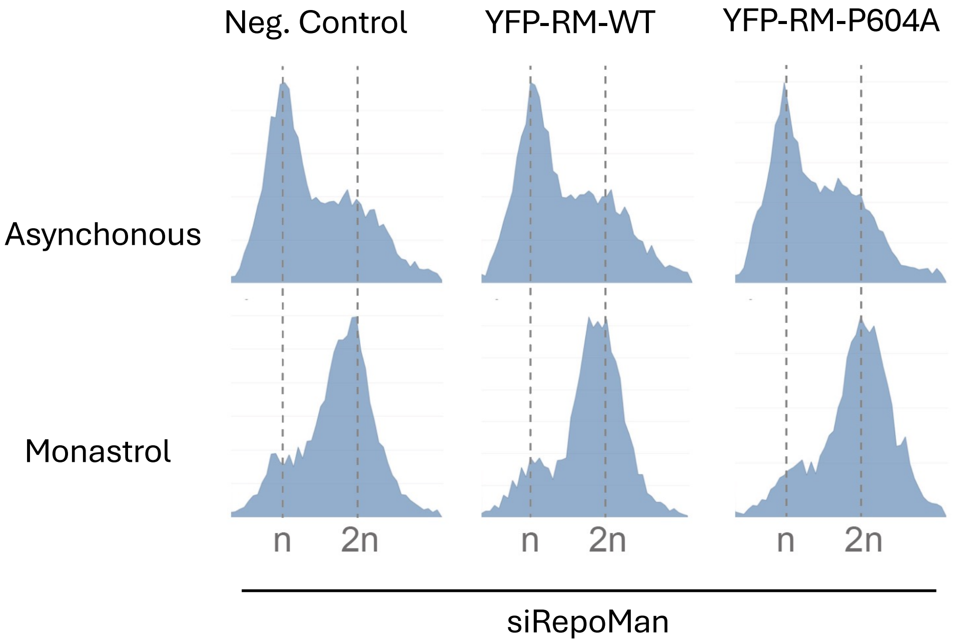

Neg. Control
YFP-RM-WT
YFP-RM-P604A
Asynchonous
Monastrol
n
2n
n
2n
n
2n
siRepoMan

Figure S5

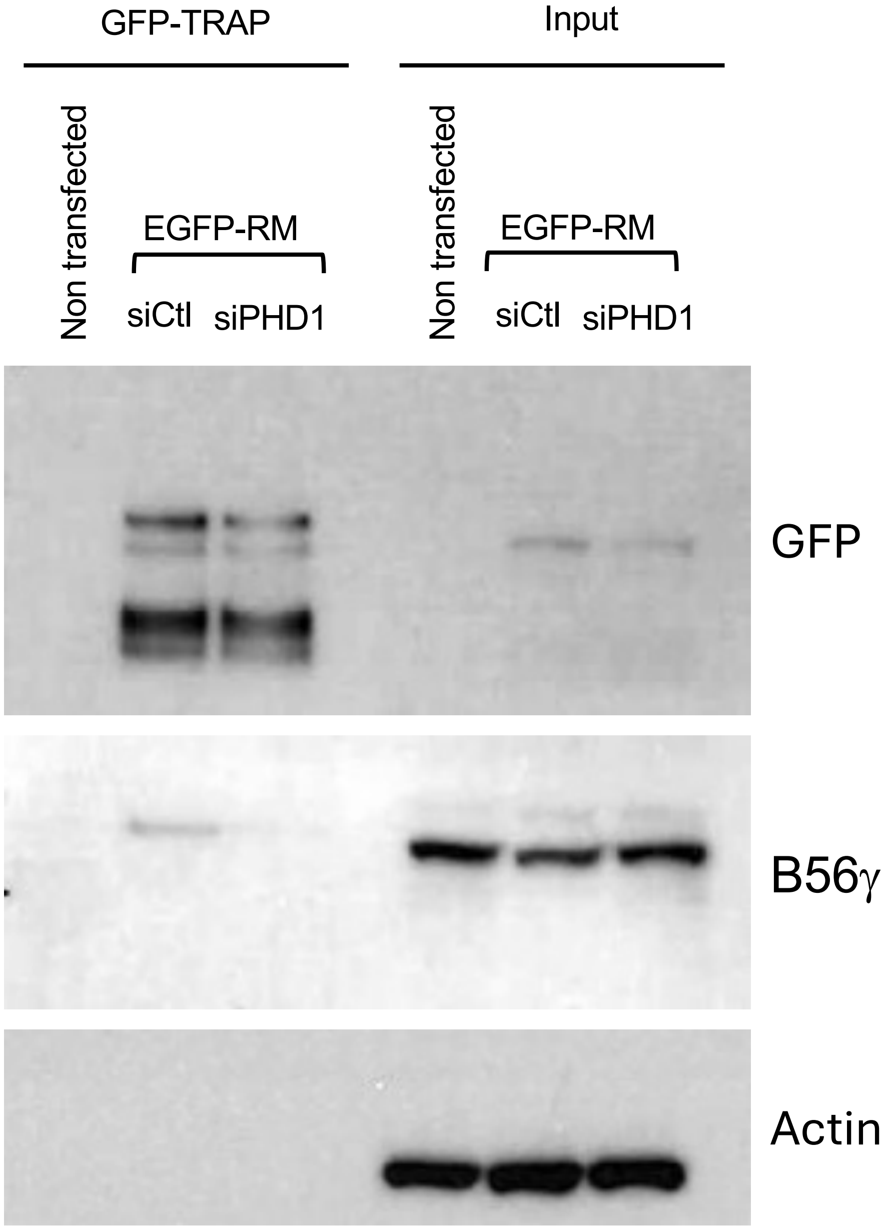

GFP-TRAP
Input
Non transfected
EGFP-RM
siCtl
siPHD1
Non transfected
EGFP-RM
siCtl
siPHD1
GFP
B56γ
Actin

### Figure S6

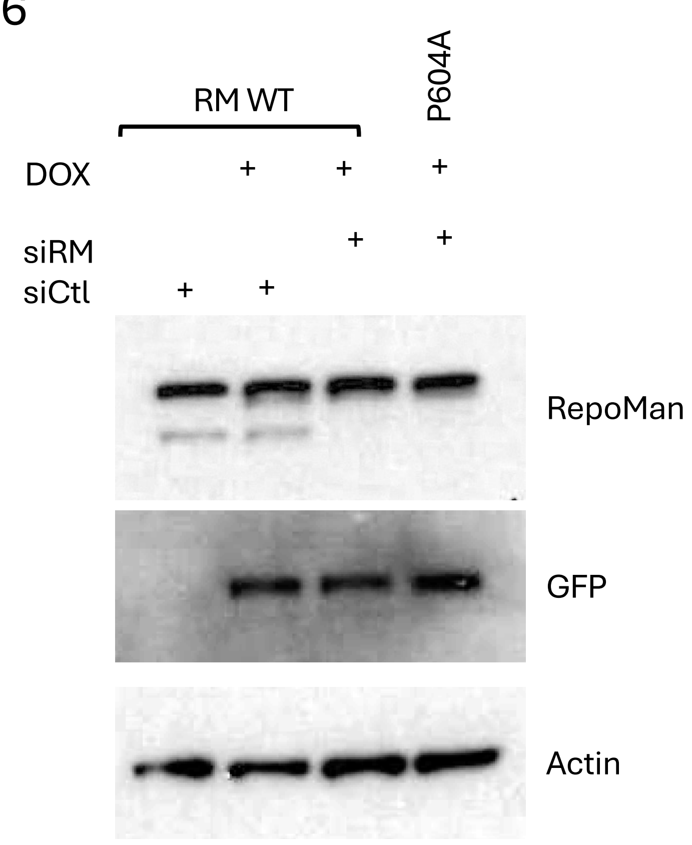
